# Computed Tomography Measures of Inter-site tumor Heterogeneity for Classifying Outcomes in High-Grade Serous Ovarian Carcinoma: a Retrospective Study

**DOI:** 10.1101/531046

**Authors:** Harini Veeraraghavan, H. Alberto Vargas, Alejandro-Jimenez Sanchez, Maura Miccó, Eralda Mema, Marinela Capanu, Junting Zheng, Yulia Lakhman, Mireia Crispin-Ortuzar, Erich Huang, Douglas A. Levine, Joseph O. Deasy, Alexandra Snyder, Martin L. Miller, James D. Brenton, Evis Sala

## Abstract

**Abstract:** *Background:* High grade serous ovarian carcinoma shows marked intra-tumoral heterogeneity which is associated with decreased survival and resistance to platinum-based chemotherapy. Pre-treatment quantification of spatial tumor heterogeneity by multiple tissue sampling is not clinically feasible. Using standard-of-care CT imaging to non-invasively quantify heterogeneity could have high clinical utility and would be highly cost-effective. Texture analysis measures local variations in computed tomography (CT) image intensity. Haralick texture methods are typically used to capture the heterogeneity of entire lesions; however, this neglects the possible presence of texture habitats within the lesion, and the differences between metastatic sites. The primary aim of this study was to develop texture analysis of intra-site and inter-site spatial heterogeneity from standard-of-care CT images and to correlate these measures with clinical and genomic features in patients with HGSOC.

*Methods and findings:* We analyzed the data from a retrospective cohort of 84 patients with HGSOC consisting of 46 patients from Memorial Sloan Kettering Cancer Center (MSKCC) and 38 non-MSKCC cases selected from The Cancer Imaging Archive (TCIA). Inclusion criteria consisted of FIGO stage II–IV HGSOC, attempted primary cytoreductive surgery, intravenous contrast-enhanced CT of abdomen and pelvis performed prior to surgery and availability of molecular tumor data analysed as per the Cancer Genome Atlas (TCGA) Research Network ovarian cancer project. Manual segmentation and image analysis was performed on 463 metastatic tumor sites from 84 patients. In the MSKCC cohort the median number of tumor sites was 7 (interquartile range 5–9) and 4 (interquartile range 3–4) in the TCIA patients. Sub-regions were produced within each tumor site by grouping voxels with similar Haralick texture using the Kernel K-means method. We derived statistical measures of intra- and inter-site tumor heterogeneity (IISTH) including cluster sites entropy (cSE), cluster sites standard deviation (cluDev) and cluster sites dissimilarity (cluDiss) from sub-regions identified within and between individual tumor sites. Unsupervised clustering was used to group patient IISTH measures into low, medium, high, and ultra-high heterogeneity clusters from each cohort. The IISTH measure cluDiss was an independent predictor of progression-free survival (PFS) in multivariable analysis in both datasets (MSKCC hazard ratio [HR] 1.04, 95% CI 1.01–1.06, *P* = 0.002; TCIA HR 1.05, 95% CI 1.00–1.10, *P* = 0.049). Low and medium IISTH clusters were associated with longer PFS in multivariable analysis (MSKCC HR 2.94, 90% CI 1.29–6.70, *P* = 0.009, TCIA HR 5.94, 95% CI 1.05–33.6, *P* = 0.044). IISTH measures were robust to differences in the CT imaging systems. Average Haralick textures contrast (TCIA HR 1.08, 95% CI 1.01–1.10, *P* = 0.019) and homogeneity (TCIA HR 1.09, 95% CI 1.02–1.16, *P* = 0.008) were associated with PFS in mutivariate analysis only in the TCIA dataset. All other average Haralick textures and total tumor volume were not associated with PFS in either dataset.

*Conclusions:* Texture measures of intra- and inter-site tumor heterogeneity from standard of care CT images are correlated with shorter PFS in HGSOC patients. These quantitative methods are independent of the CT imaging system and can thus be applied in clinical practice. The methodology proposed here enables the non-invasive quantification of intra-tumoral heterogeneity and disease stratification for future experimental medicine studies and clinical trials, particularly in cases where total tumour volume and averaged textures have low predictive power.

**Author summary:** *Why was this study done?:*

- Tumor heterogeneity is a feature of many solid malignancies including ovarian cancer.
- Recent genomic research suggests that intra-site tumor heterogeneity (heterogeneity within a single tumor site) and inter-site tumor heterogeneity (heterogeneity between different metastatic sites in the same patient) correlate with clinical outcome in HGSOC.

*What did the researchers do and find?:*

- We developed quantitative and non-invasive image-analysis based measures for predicting outcome in HGSOC patients by combining image-based information from within and between multiple tumor sites.
- Using datasets from two sources, we demonstrate that these image-based tumor heterogeneity measures predict progression free survival in patients with HGSOC.

*What do these findings mean?:*

- Non-invasive measures of CT image heterogeneity may predict outcomes in HGSOC patients.
- Wider application of these CT image heterogeneity measures could prove useful for stratifying patients to different therapies given that total tumour volume and averaged textures have low predictive power.

## Introduction

High grade serous ovarian carcinoma (HGSOC) is the deadliest gynecologic malignancy with overall survival remaining unchanged over the last 20 years [1]. Although HGSOC shows marked sensitivity to platinum-based chemotherapy [2], the majority of cases recur and become progressively resistant to subsequent treatment regimes [3]. Acquisition of resistance may be related to specific mutational processes that drive genomic heterogeneity [4, 5] and clonal evolution [6, 7]. HGSOC exhibits marked intra-site and inter-site genetic heterogeneity across metastatic sites in the peritoneal cavity [5–7] with altered immunological infiltrates and tumor microenvironments [8]. Detection of spatial or temporal heterogeneity by multiple sampling in a single patient is expensive, invasive, and often clinically impractical. Consequently, analysis of heterogeneity has only been performed on a limited number of patients with HGSOC [5–7].

Computed tomography (CT) and serum CA-125 measurement are routinely used for initial staging and treatment monitoring of patients with HGSOC, but standard assessments do not provide information on tumor heterogeneity. Texture analysis of CT data is a radiomics method [9–11] that can provide detailed quantitative characterization of local variations in intensity levels throughout an image. Texture measures are computed from regions of interest (ROI) drawn to delineate tumors on cross-sectional images. The local texture can be measured using Haralick methods that quantify the second order intensity variation within an ROI and other high-order features based on grey-level matrices [12]. Radiomics methods have been applied to evaluate intra-tumor heterogeneity based on a single site of disease per patient in primary tumors [9, 13–16] and metastatic disease [17–19]. However, a summary CT texture value for a single tumor site does not capture the potential variability between different regions within the tumor and between multiple tumor sites in the same patient. CT-based textures that combine intra- and inter-site spatial heterogeneity have not been developed and are needed to investigate how the spatial heterogeneity between tumor sites is related to variable response and the development of resistance [20].

The ability to predict patients with poor outcomes using non-invasive imaging-based methods that quantify heterogeneity within (intra-site) and between separate tumor sites (inter-site) would have high clinical utility. Therefore the aims of this retrospective study were: (1) to develop computational texture analyses methods from standard of care CT images that incorporate both intra-site and inter-site spatial heterogeneity and (2) to correlate the texture measures with adverse prognostic genomic features, tumor immune microenvironment (TME) and outcome in patients with HGSOC evaluated as per the Cancer Genome Atlas (TCGA) Research Network ovarian cancer pilot project.

## Methods

### Ethics and consent

This study was compliant with the Health Insurance Portability and Accountability Act and received approval by the Institutional Review Board at Memorial Sloan Kettering Cancer Center with a waiver for requiring informed consent. The TCIA is a managed open-source archive of radiology images of cancer contributed by 28 different institutions that provides datasets obtained after prospective informed consent and de-identification of all protected patient health information [21].

### Study design and patients

This manuscript was written in accordance to the REMARK [22] and STROBE guidelines [23] (see STROBE checklist S1 Checklist and S1 Text for summary of statistical analysis). The study population consisted of two cohorts of patients with HGSOC: a single institution dataset from MSKCC (*n* = 46) and a multi-institution dataset (*n* = 38) from the ovarian-TCIA [24] which included patients treated at five different institutions (Fig 1). The eligibility criteria included: (i) FIGO stage II–IV HGSOC, (ii) attempted primary cytoreductive surgery, (iii) standard of care intravenous contrast-enhanced CT of the abdomen and pelvis performed prior to surgery, (iv) at least two tumor sites identified on CT in order to evaluate inter-site tumor heterogeneity, and (v) molecular analysis performed as per the The Cancer Genome Atlas (TCGA) Research Network ovarian cancer pilot project. Patients who received neoadjuvant chemotherapy prior to surgery were excluded from the study. Forty-six patients from MSKCC and 38 patients from TCIA datasets fulfilled the inclusion criteria. S1 Fig shows the REMARK diagram for selection of patients.

**Fig 1.**
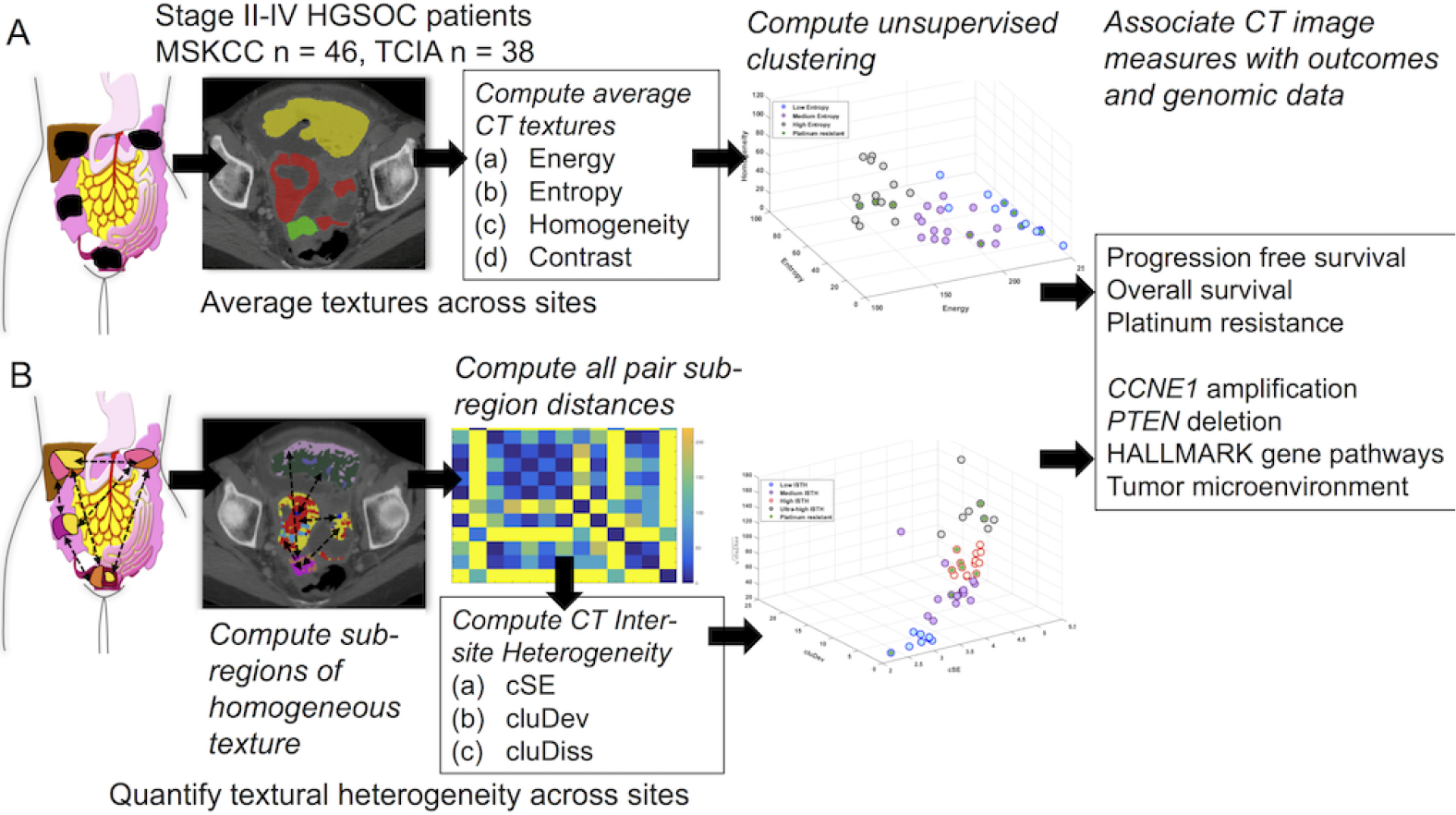
Schema of experimental workflow. For each patient, average texture heterogeneity and inter-site tumor heterogeneity texture measures were computed. Texture measures from the two cohorts of patients were used to group patients using unsupervised clustering which were correlated with outcomes.

All patients from the current study were included in two prior studies that investigated the associations between qualitative CT imaging features, Classification of Ovarian Cancer (CLOVAR) transcriptomic profiles and survival in a single and multiple institution datasets, respectively [25, 26]. Previously [27], we also evaluated the feasibility of CT-based texture measures of inter-site tumor heterogeneity in 38 patients from the MSKCC dataset. The current study now quantifies both intra-site and inter-site tumor heterogeneity and evaluates these new texture measures on an additional independent dataset. Clinical details including disease stage, residual disease after debulking surgery and platinum sensitivity were abstracted from the patient clinical records by radiologists and oncology imaging fellows from MSKCC and the institutions contributing to the TCIA dataset. Platinum resistance was defined as a platinum-free interval of less than 6 months after initial therapy [28].

We performed an exploratory analysis to identify whether the novel image-based heterogeneity measures were associated with adverse genomic factors including putative copy-number alterations to *CCNE1*, *PTEN*, *RB1*, *NF1*, somatic point mutations to *BRCA1*, and *BRCA2*, the Hallmark gene set enriched pathways, and the tumor microenvironment consisting of stromal and immune scores. The copy number alterations and point mutations were computed using the GISTIC method [29] and obtained from the cbioportal [30]. The Hallmark gene set enrichments were extracted through the analysis of the bulk tumor RNA sequence data was performed using single sample gene-set enrichment analysis [31], while the ESTIMATE method [32] was used to quantify the tumor microenvironment (TME) cell types consisting of the stromal and immune signatures from the RNA sequence data.

### Segmentation of CT images and extraction of texture features

Two oncologic imaging research fellows with 4 and 6 years experience respectively evaluated all CTs and manually segmented primary and suspected metastatic tumor sites in the abdomen and pelvis. Manual segmentation was performed using 3DSlicer [33] by tracing the tumor contour on each CT slice to produce a volume of interest (VOI) for each lesion. The CT images were rescaled from 0–255 prior to computation of the texture measures. Texture analysis was performed using in-house software implemented in C++ using the Insight ToolKit (ITK) [34]. Voxel-wise Haralick textures [12] were computed within each manually delineated VOI using a neighborhood size of 5 × 5 × 1 and thirty two histogram bins for the grey level co-occurrence matrix.

### Average heterogeneity of tumor burden

Average heterogeneity of all tumor sites was derived as the mean of voxel-wise Haralick texture values from all sites within each patient. Haralick energy, entropy, homogeneity and contrast were computed for the primary lesion and every metastatic site. Tumor volume was estimated as the total number of voxels within each VOI multiplied by voxel size.

### Intra and inter-site tumor heterogeneity

The heterogeneity of the disease was quantified according to the following procedures (see also S1 Methods):

(i) **Division of lesions into distinct sub-regions.** First, voxel-wise textures were derived by sliding a fixed size patch (5 × 5) across the whole image (texture measures within a patch were assigned to the central voxel within that patch). Then, sub-regions of homogeneous texture were extracted within each tumor by grouping voxels with similar textural values (Haralick textures) using kernel K-means clustering [35] (Fig 2A). The number of clusters was set to a maximum of five and the best number for each patient was determined using Akaike information criterion. Each sub-region was described by the collection of mean values of the four individual Haralick texture measures. These sub-regions are the primary input to the heterogeneity metrics.
(ii) **Quantification of the similarity between all pairs of sub-regions.** Textural dissimilarity between all pairs of sub-regions was computed as the Euclidean distance of the sub-region textures. As a result, sub-regions with highly distinct textures would result in very large dissimilarities. All pairs of sub-region dissimilarities were summarized as a dissimilarity map, and discretized into 10 bins(Fig 2A).
(iii) **Definition of patient-level heterogeneity metrics. Two types of heterogeneity metrics were defined based on the discretized dissimilarity values.**

1. Frequency-based metric. The frequencies of the pairwise sub-region dissimilarities were normalized in the range of 0-1 for each patient and discretized into ten bins and used to compute the cluster sites entropy (cSE), that measures the entropy in clusters of uniform textures or sub-regions computed within the tumor sites defined and using the Shannon entropy formulation. cSE is a measure of two components, namely the number of unique dissimilarities (or diversity) and their equality in the distribution (or evenness) across all tumor sites.
2. 2D histogram-based metrics. A new class of features was designed in order to capture the diversity of distinct groups of texturally similar sub-regions. To this end, the dissimilarity map was used to create a new 2D histogram called the ‘group size dissimilarity matrix’ (GDM), that captured how many pairs of sub-regions have similar levels of dissimilarity. From this matrix, two scalar metrics were extracted, called cluDev and cluDiss. The cluDev measure is the standard deviation of the GDM. The cluDiss measure quantifies the richness of distinct groups of subregions (grouped by their even dissimilarities) and the spread of the group dissimilarities. This measure adds to the cSE metric by magnifying the asymmetry in the distribution of dissimilarities. Whereas the cSE measure focuses on diversity defined as a function of number of unique dissimilarities and their frequencies, the cluDiss measure focuses on the relatedness between groups of subregions when modeling the diversity of dissimilarities. For example, tumor sites containing sub-regions with highly similar textures will produce low cluDiss value and low cSE or low heterogeneity (Fig 2B). Conversely, the presence of highly distinct sub-regions across the tumor sites will produce a high cluDiss value and higher cSE (Fig 2C) corresponding to high intra- and inter-tumor heterogeneity. The main difference between the two measures is that cSE has a restricted range and only quantifies dissimilarities while the cluDiss focuses on amplifying the extent of dissimilarities. All of the three measures are correlated to the number of metastatic sites S5 Table and are strongly correlated with one another S1 Fig. Note that all sub-regions, regardless of their site of origin, are treated on the same footing. The metrics defined above are therefore referred to as intra- and inter-site tumor heterogeneity (IISTH) features, as they capture both levels simultaneously. The source code for generating the IISTH features is available for download from (Github link).
(iv) **Unification of metrics into single IISTH score.** All three measures of inter-site tumor heterogeneity show strong correlation with each other (S1 Fig). Therefore, the three measures were combined to produce a combined intra/inter-site tumor heterogeneity score (IISTH). The combined score or IISTH score was computed through factor analysis (available through the psych package in R) of the measures and using the factor loadings as the combined score for correlating with the survival and outcomes.

The mutual complementarity of the 3 IISTH measures was explored in two different ways. Firstly, the three measures were used to cluster patients into four (as determined automatically by algorithm) different categories of increasing heterogeneity. This method has the advantage of taking into account subtle differences between the three features at the time of classifying the patients. Secondly, the three features were combined into a single score, denoted the “composite IISTH score”. This method has the advantage of providing a single heterogeneity scale along which patients can be mapped.

**Fig 2.**
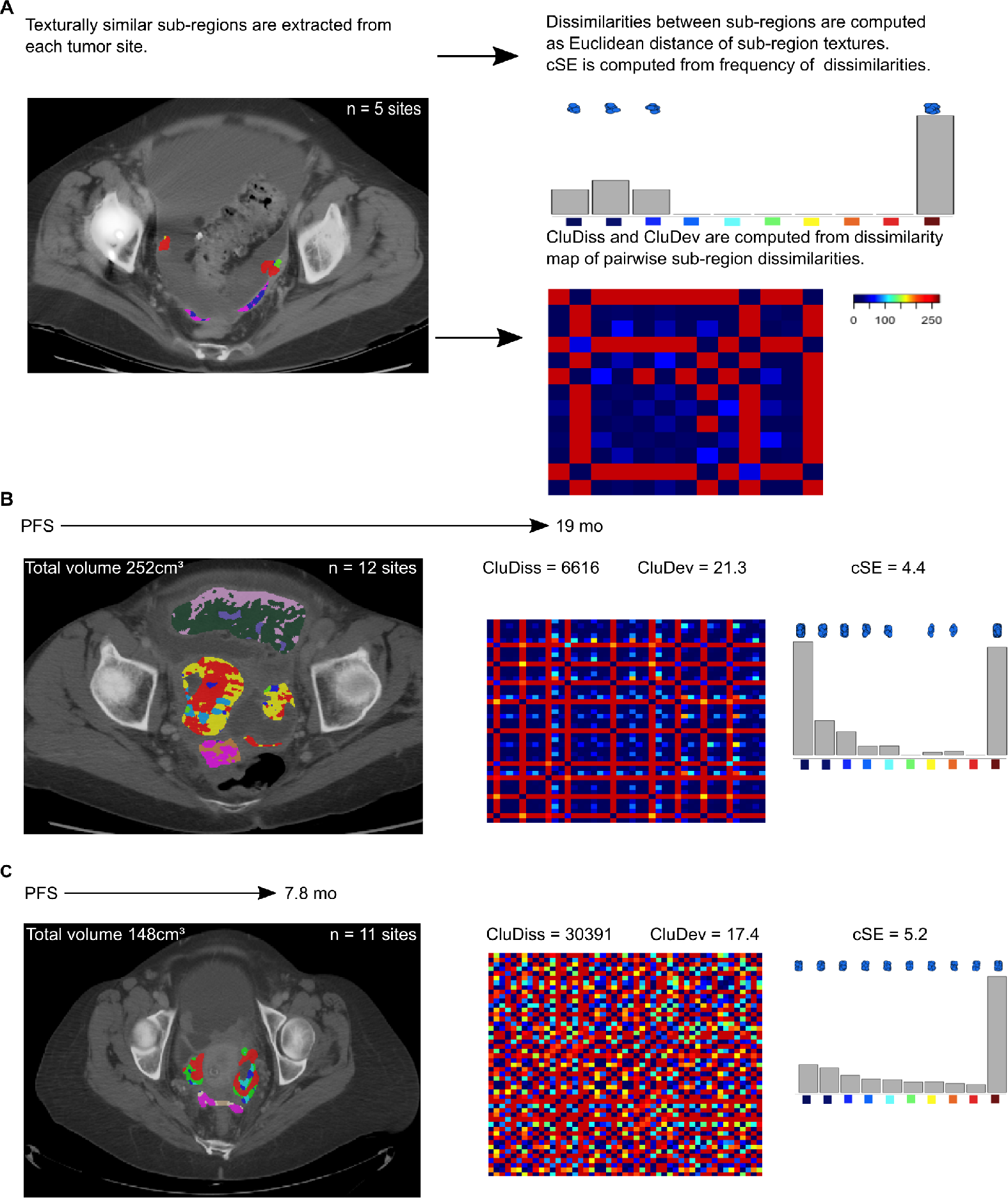
IISTH measures in 2 representative patients. (A) Method for extracting inter-site tumor heterogeneity measures. Sub-regions with homogeneous textures are extracted for each tumor location. Sub-regions with different textures are indicated by different colored regions in the CT image. The dissimilarity between sub-regions are computed using Euclidean distances of the sub-region textures. Cluster sites entropy (cSE) is computed from the distribution of dissimilarities. The cluDiss and cluDev measures are computed from a dissimilarity map that expresses the pairwise sub-region dissimilarities. (B) Patient with smaller IISTH measures and associated with better outcomes (PFS of 19 mo) and (b). Patient with larger IISTH measures and associated with worse outcomes (PFS of 7.8 mo).

### Unsupervised clustering of patients by heterogeneity measures

Self-tuning spectral clustering [36] was used to group patients by their heterogeneity into distinct clusters separately using the proposed IISTH and the average tumor heterogeneity measures. Self-tuning spectral clustering automatically determines the appropriate number of clusters required to partition the data by minimizing the cost of data splits. The relative proportion of the various outcome measures in each cluster was then computed.

### Hallmark gene sets estimation

Pathway enrichment analysis was performed to extract the Hallmark gene sets for each sample from the MSigDB database version 6.1 [37]. Single-sample gene set enrichment analysis [31], a modification of standard gene set enrichment analysis (GSEA) [38], was performed on RNA measurements for each sample using the GSVA package version 1.28.0 [39] in R version 3.5.0 using the ssgsea method and tau = 0.25. Next, gene sets for each cell type were generated using the union of genes derived from the different methods. Thus, the cell types were represented by the union of genes of the different methods. Normalized enrichment scores were generated for the computed Hallmark gene sets [40]. ESTIMATE method [32] was used to quantify the immune and stromal signatures from RNA-seq data.

### Statistical analysis

Pre-specified analyses were determined after data collection but prior to performing statistical analysis. Additional exploratory analysis, namely, determining the association of texture measures to genetic alterations was performed after the extraction of texture-based clusters. Patient characteristics and texture measures were summarized using standard descriptive statistics using median and interquartile range (IQR). Cox regression was used to assess associations between OS and PFS and texture measures after adjusting for age, disease stage, resection status, and tumor volume. Cox analysis for tumor volume was performed after adjusting for age and disease stage. Kaplan-Meier curves were used to estimate OS and PFS. Associations between categorical variables, including platinum-resistance status and genetic alternations in *CCNE1* in the texture-based clusters were examined using Fisher’s exact test. Comparison between different CT scanner data, including Haralick texture and IISTH measures, was studied using the Wilcoxon rank sum test. Data with missing variables were excluded from the analysis. Spearman rank correlation was performed to assess the correlation between the continuous texture measures and tumor volume, number of tumor sites and molecular factors. Associations were investigated separately for each institution. Values of *P* < 0.05 were considered to be significant. Owing to limited hypothesis testing and the exploratory nature of the study, we did not adjust for multiple testing. Thus, the reported results are preliminary and require further validation in future investigations.

## Results

### Patient characteristics

A total of 9625 CT images were collected from 84 patients with HGSOC. The median number of tumor sites analysed was 7 (IQR 5–9) for the MSKCC cohort and 4 (IQR 3–4) for the TCIA cohort. In total, 463 volumes of interest (VOI) were analysed to obtain intra- and inter-site tumor heterogeneity (IISTH) texture measures for 84 patients using computational methods summarized in S1 Methods.Fig 1 summarizes the experimental design. S2 Fig shows the REMARK diagram for selection of patients.

Patient characteristics for the two datasets are summarized in Table 1. The median follow-up was 41 months (IQR 24–55 mo) in the MSKCC cohort and 21 months (IQR 7–39 mo) in the TCIA cohort, with all but 2 patients in MSKCC and 19 in TCIA progressing during the follow-up period. Comparison of the MSKCC and TCIA datasets showed significant differences between the two datasets for progression-free survival (PFS), CT-derived total tumor volume and for texture measures (Table 1). Owing to these differences, it was not possible to use the two datasets in a training–testing–validation analysis. We therefore analyzed the two datasets separately and assessed whether the same trends were observed for the association of texture measures and outcomes.

**Table 1.** Retrospective cohort characteristics.

| Patient characteristic | MSKCC | TCIA |
| --- | --- | --- |
| Number of patients | 46 | 38 |
| Age (median)(IQR) | 59 (50–67) | 62 (53–72) |
| Stage at diagnosis (proportion of patients) |  |  |
| II | 0 (0%) | 4 (11%) |
| III | 31 (67%) | 29 (76%) |
| IV | 15 (33%) | 5 (13%) |
| Surgical debulking outcome (number of patients) |  |  |
| Complete | 15 (33%) | 8 (21%) |
| Optimal $\leq 1$ cm | 23 (50%) | 16 (42%) |
| Suboptimal $> 1$ cm | 8 (17%) | 12 (32%) |
| Unknown | 0 (0%) | 2 (5%) |
| Recurrence status* (number of patients) |  |  |
| Recurring | 44 (96%) | 19 (50%) |
| Not recurring | 2 (4%) | 19 (50%) |
| Disease status* (number of patients) |  |  |
| Alive | 17 (37%) | 16 (42%) |
| Dead | 29 (63%) | 22 (58%) |
| Follow up* mos (median)(IQR) | 41 (24–55) | 21 (7–39) |
| Survival (median)(IQR) |  |  |
| PFS<sup>†</sup> mos | 16 (10–27) | 17 (8–25) |
| OS<sup>†</sup> mos | 59 (45–77) | 31 (16–50) |
| Platinum status (number of patients) |  |  |
| Sensitive | 35 (76%) | 17 (45%) |
| Resistant | 9 (20%) | 7 (18%) |
| Unknown | 2 (4%) | 14 (37%) |
| tumor volume* (cm<sup>3</sup>)(median)(IQR) | 122 (65–232) | 328 (153–601) |
| tumor sites* (median)(IQR) | 7 (5–9) | 4 (3–4) |

\* indicates datasets were significantly different ( $P < 0.05$ ).
<sup>†</sup> indicates datasets were significant different ( $P < 0.05$ ) computed using Log-rank tests. The reported number of events occurred within the time frame of the study.

### Intra- and inter-site tumor heterogeneity CT texture-based measures predict outcome

Unsupervised clustering using the IISTH measures resulted in four clusters with low, medium, high and ultra-high IISTH heterogeneity (Fig 3). Unsupervised clustering of average texture heterogeneity resulted in three clusters with low, medium, and high texture entropy (S3 Fig).

**Fig 3.**
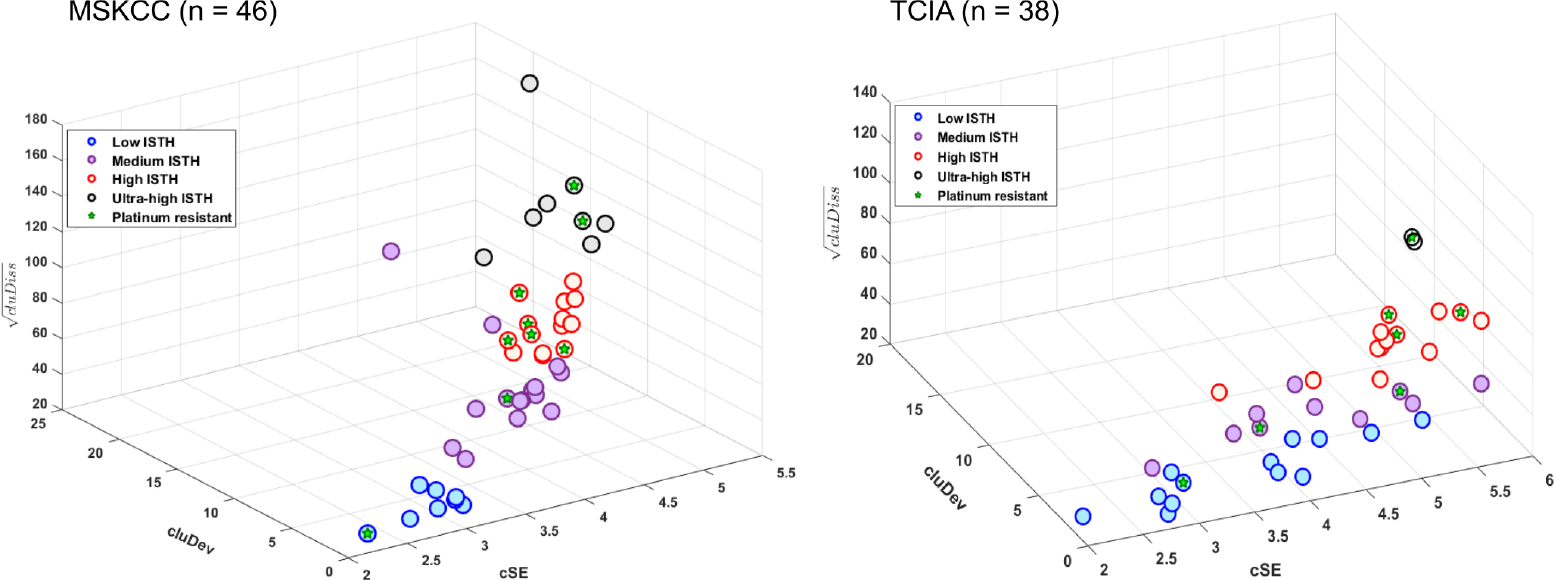
Unsupervised clustering of MSKCC and TCIA cohorts using IISTH measures.

In the univariable analysis, the IISTH measures cSE, cluDiss, composite IISTH scores, and the IISTH clusters were all significant predictors of PFS (Table 2) in both MSKCC and TCIA datasets. When adjusted for age, disease stage, tumor volume and surgical resection status, only cluDiss remained significant in both datasets (MSKCC hazard ratio [HR] 1.10, 95% CI 1.02–1.20, *P* = 0.015, TCIA HR 1.40, 95% CI 1.15–1.80, *P* = 0.001) (Figure 4). Multivariable analysis of the composite heterogeneity measure (or IISTH score) showed that the same measure was a significant predictor of outcome (MSKCC HR 1.51, 95% CI 1.01–2.26, *P* = 0.43, TCIA HR 10.00, 95% CI 2.62–38.12, *P* = 0.0007) (SI Fig, S4 Fig). Multivariable analysis also showed that the IISTH clusters were significant predictors of outcome (MSKCC HR 2.58, 95% CI 1.23–5.40, *P* = 0.012, TCIA HR 10.74, 95% CI 2.67–43.2, *P* = 0.001) Fig 5).

**Table 2.** Univariable (unadjusted) and multivariable (adjusted) analysis of pre-treatment values as predictors for progression free survival (PFS) for MSKCC and TCIA dataset. Multivariable Cox hazard regression was adjusted for age, stage, resection status, and volume.

| Variable | MSKCC |  |  |  |  |  | TCIA |  |  |  |  |  |
| --- | --- | --- | --- | --- | --- | --- | --- | --- | --- | --- | --- | --- |
|  | Univariable |  |  | Multivariable |  |  | Univariable |  |  | Multivariable |  |  |
|  | HR | CI | P-Value | HR | CI | P-Value | HR | CI | P-Value | HR | CI | P-Value |
| cSE | 1.60 | 1.10–2.40 | 0.027 | 2.36 | 1.11–5.00 | 0.026 | 3.00 | 1.30–6.80 | 0.007 | 2.74 | 0.47–15.8 | 0.26 |
| cluDiss | 1.10 | 1.02–1.20 | 0.001 | 1.04 | 1.01–1.06 | 0.002 | 1.20 | 1.10–1.40 | 0.001 | 1.05 | 1.00–1.10 | 0.049 |
| cluDev | 1.00 | 0.98–1.10 | 0.140 | 1.05 | 0.95–1.20 | 0.338 | 1.60 | 1.09–2.30 | 0.016 | 0.86 | 0.36–2.00 | 0.731 |
| IISTH score | 1.63 | 1.13–2.34 | 0.009 | 3.20 | 1.45–7.08 | 0.004 | 2.16 | 1.26–3.72 | 0.005 | 2.25 | 0.62–8.23 | 0.218 |
| IISTH cluster (3,4/1,2)† | 2.77 | 1.39–5.50 | 0.004 | 2.94 | 1.29–6.70 | 0.009 | 6.85 | 2.27–20.7 | < 0.001 | 5.94 | 1.05–33.6 | 0.044 |
| Energy | 1.00 | 0.99–1.01 | 0.912 | 0.99 | 0.98–1.00 | 0.066 | 0.99 | 0.98–1.00 | 0.228 | 0.99 | 0.97–1.00 | 0.224 |
| Entropy | 0.99 | 0.97–1.01 | 0.208 | 1.01 | 1.00–1.03 | 0.056 | 1.01 | 0.99–1.03 | 0.196 | 1.03 | 0.99–1.10 | 0.09 |
| Contrast | 0.99 | 0.98–1.00 | 0.204 | 0.99 | 0.98–1.00 | 0.159 | 1.03 | 0.97–1.08 | 0.341 | 1.08 | 1.01–1.10 | 0.019 |
| Homogeneity | 0.99 | 0.98–1.01 | 0.815 | 1.01 | 0.99–1.03 | 0.552 | 1.02 | 0.99–1.04 | 0.247 | 1.09 | 1.02–1.16 | 0.008 |
| Haralick cluster (3/1)† | 1.20 | 0.57–2.55 | 0.634 | 1.88 | 0.79–4.50 | 0.155 | 2.40 | 0.67–8.62 | 0.179 | 1.62 | 0.31–8.50 | 0.57 |
| Haralick cluster (2/1)† | 1.17 | 0.55–2.51 | 0.678 | 1.57 | 0.70–3.50 | 0.271 | 1.60 | 0.31–8.20 | 0.576 | 0.96 | 0.16–5.80 | 0.967 |
| Resection (sub-optimal) | 3.70 | 1.44–9.50 | 0.007 | 2.97 | 0.82–10.70 | 0.097 | 0.95 | 0.25–3.60 | 0.946 | 1.45 | 0.28–7.40 | 0.657 |
| Resection (optimal) | 1.71 | 0.85–3.40 | 0.130 | 1.33 | 0.49–3.60 | 0.575 | 0.98 | 0.33–2.90 | 0.973 | 1.13 | 0.279–4.60 | 0.860 |
| Stage (IV/III,II) | 1.10 | 0.59–2.10 | 0.730 | 1.15 | 0.53–2.50 | 0.715 | 1.40 | 0.41–4.90 | 0.58 | 1.12 | 0.29–4.40 | 0.872 |
| Age (years) | 1.10 | 1.02–1.10 | 0.003 | 1.05 | 1.01–1.10 | 0.010 | 1.00 | 0.97–1.00 | 0.796 | 1.06 | 0.99–1.10 | 0.056 |
| Volume (cm<sup>3</sup>) | 1.00 | 1.00–1.00 | 0.530 | 1.00 | 0.99–1.00 | 0.968 | 1.00 | 1.00–1.00 | 0.320 | 1.00 | 0.99–1.00 | 0.742 |
| Sites | 1.10 | 1.00–1.30 | 0.028 | 1.02 | 0.86–1.20 | 0.830 | 1.60 | 1.10–2.40 | 0.012 | 2.44 | 1.36–4.40 | 0.003 |

† 3, 4 are high and ultra-high heterogeneity clusters and 1, 2 are low and medium heterogeneity clusters.
For continuous measures, HR > 1, an increase in the value is associated with higher risk or number of events and decreased PFS. For binary variables, the HR listed is the first option and HR=1 is the second option. HR < 1 indicates lowering of risk with the same option.
CI - confidence interval HR - hazard ratio.
Cox analysis for cluDiss as a continuous measure was computed after computing square root of cluDiss to achieve reasonable data scaling.

**Fig 4.**
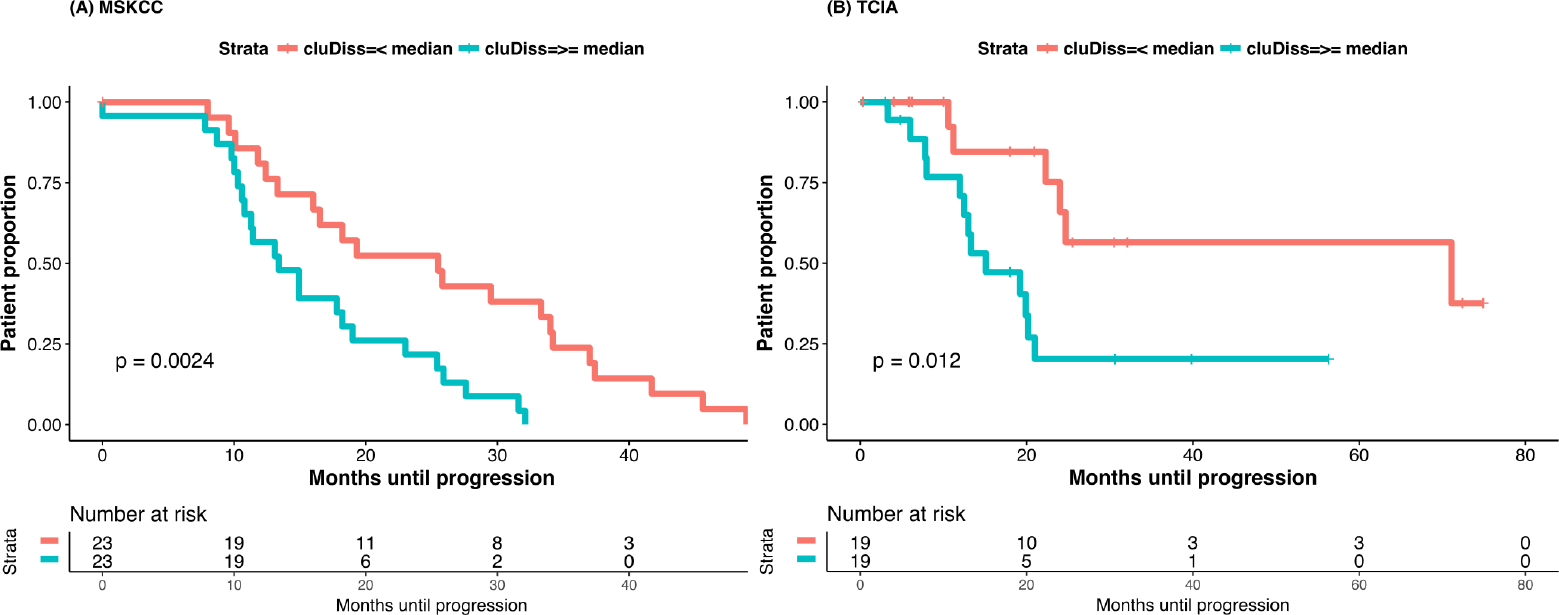
Survival by cluDiss measures in MSKCC and TCIA cohorts Kaplan-Meier analysis using groups split on median cluDiss value (MSKCC = 6529; TCIA = 2896).

**Fig 5.**
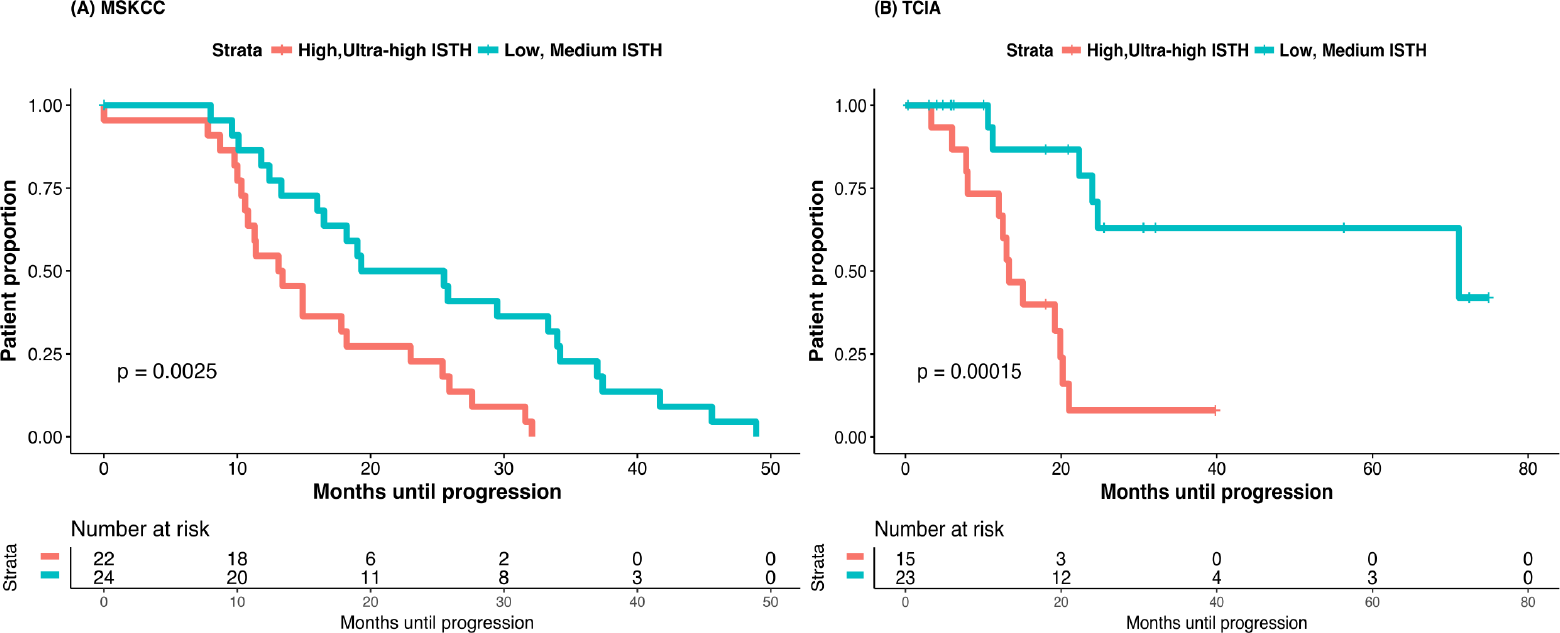
Survival by IISTH clusters (low, medium IISTH versus high, ultra-high IISTH) obtained through self-tuning spectral clustering for MSKCC and TCIA datasets.

Although sub-optimal resection and age were independent predictors of PFS in the multivariable analysis in the MSKCC dataset, neither measure was associated with PFS in the TCIA dataset. Similarly, tumor volume did not predict PFS.

The IISTH measure cluDiss, IISTH cluster, and Haralick homogeneity texture were significant predictors of OS (S2 Table) in the univariable analysis for the MSKCC dataset. When adjusted for age, disease stage, tumor volume, number of metastatic sites, and surgical resection status, using the Cox proportional hazards model, IISTH cluster remained significant predictor of OS in the MSKCC dataset.

The IISTH measures were not correlated with total tumor volume but showed strong positive correlation with the number of tumor sites (S5 Table). Average Haralick tumor heterogeneity measures showed variable (some positive and others negative) correlation with tumor volume (S5 Table).

The high and ultra-high IISTH heterogeneity clusters in the MSKCC dataset had a larger number of patients with primary platinum resistance (7 versus 2) compared to low and medium heterogeneity clusters (S3 Table). However there was no significant difference in the prevalence of platinum resistance (MSKCC *P* = 0.064, TCIA *P* = 1.0) between the two cluster groups. Similar to the IISTH clusters, clusters of average tumor heterogeneity (S4 Table) did not show an association with platinum resistance (MSKCC *P* = 0.73, TCIA *P* = 0.57).

### IISTH cluDiss measure of pelvic metastatic disease is negatively correlated with Hallmark gene sets

Given that only the cluDiss measure was associated with PFS in the multivariate analysis, we evaluated whether cluDiss correlated with Hallmark gene set enrichment scores. We computed the IISTH cluDiss measure for the pelvic (ovarian mass and cul-de-sac) metastases to assess its association to the genomic pathways estimated using ssGSEA [40]. We focused on the pelvic metastatic disease for the purpose of this analysis as the tissue samples for molecular analysis were obtained from the pelvic disease. The cluDiss measure was normalized for the data to have zero mean and unit standard deviation using z-score standardization and to facilitate easier visualization of the correlations. The cluDiss measure had a negative correlation with WNT enrichment (*ρ* = −0.35; *P* = 0.028) (Fig 6). The WNT/Beta-Catenin signaling enrichment was highest in the IISTH cluster with the least heterogeneity and decreased in IISTH clusters with increasing texture heterogeneity. Prior work by our group has shown that enrichment of WNT pathway is negatively correlated with immune-infiltration [41].

**Fig 6.**
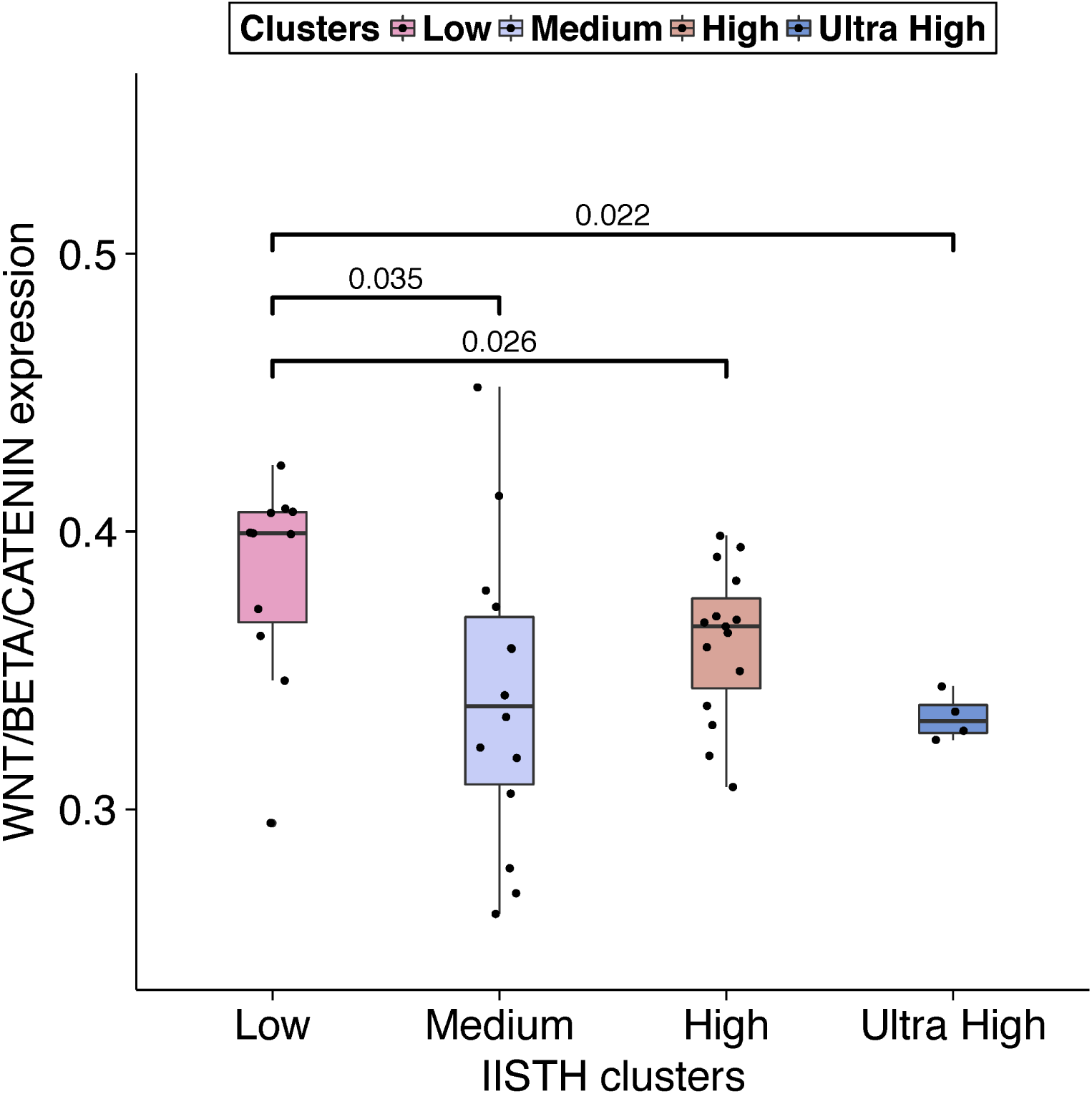
Distribution of the WNT/Beta-Catenin signaling expression across IISTH clusters of increasing heterogeneity.

### Prevalence of genetic copy number alterations in the texture heterogeneity clusters

We explored whether the IISTH clusters were associated with prognostic copy number alterations in known oncogenes and tumor suppressor genes associated with HGSOC. S5 Fig shows association of IISTH-derived clusters using patients from both datasets with alterations in *CCNE1*, *PTEN*, *RB1*, *NF1* and somatic mutations in *BRCA1* and *BRCA2*. The relative prevalence computed as the ratio of the number of patients with and without *CCNE1* amplifications was the highest in the ultra-high cluster and the lowest in the low heterogeneity cluster but there was no significant difference in the prevalence of *CCNE1* amplifications between the two cluster groups (MSKCC *P* = 0.31, TCIA *P* = 0.72). None of the patients with mutation to *BRCA1* occurred in the low heterogeneity cluster while four out of the seven with amplification or mutations to *BRCA2* occurred in the low heterogeneity cluster.

### IISTH measures were robust to differences in CT scanner manufacturer

All the MSKCC scans were obtained using General Electric (GE) CT scanners. The CT scans from the TCIA cohort were obtained from multiple institutions using different CT scanner manufacturers. We therefore investigated whether the IISTH measures were dependent on the scanner manufacturer. Wilcoxon rank sum test showed no significant difference in the distribution of IISTH texture measures between non-GE (*n* = 14; Siemens 12, Toshiba 1, Philips 1) and GE (*n* = 24) scanners (Table 3). By contrast, three out of the four Haralick textures showed significant differences between the scanners. For example, the Haralick energy measures were significantly lower (*P* = 0.001) in GE compared to non-GE scanners. Conversely, entropy was significantly higher in GE scanners compared to non-GE scanners (*P* = 0.0006). These results suggest that the IISTH measures are not influenced by CT scanner manufacturers and are potentially more applicable to multi-institutional studies than standard Haralick textures.

**Table 3.** Differences in texture measures between scanner manufacturers from TCIA dataset.

| Measure | GE |  | non-GE |  | P-Value |
| --- | --- | --- | --- | --- | --- |
|  | Median | IQR | Median | IQR |  |
| cSE | 3.2 | 2.7–3.9 | 3.2 | 3.0–3.7 | 0.56 |
| cluDev | 1.2 | 0.78–2.7 | 1.7 | 1.02–3.0 | 0.26 |
| cluDiss | 3049 | 1240–4841 | 2896 | 1452–4434 | 1.00 |
| Energy | 137 | 124–166 | 183 | 173–198 | 0.001 |
| Entropy | 72 | 56–85 | 39 | 36–52 | < 0.001 |
| Homogeneity | 11.9 | 9.9–22 | 7.8 | 4.8–13 | 0.034 |
| Contrast | 2.15 | 1.48–3.65 | 1.90 | 1.00–3.17 | 0.43 |

## Discussion

Intratumoral heterogeneity may be a critical determinant of outcome in HGSOC. We developed standard of care CT-based texture measures that model both intra- and inter-site tumor heterogeneity using IISTH measures. These capture the variation in dissimilarities within and between tumor sites and quantify the textural differences between tumour sub-regions. Our study is part of the NCI initiative to combine molecular analysis with case-matched CT images from the TCIA ovarian cancer projects [24] and therefore is representative of patients from multiple institutions. We focused on a small set of texture measures (cSE, cluDev, and cluDiss) to eliminate any false discoveries owing to seemingly promising correlations arising from a small dataset.

The IISTH measures predict progression-free survival in two different datasets. Patients with lower values of cSE and cluDiss had significantly longer PFS independently of other variables. Importantly, tumor volume did not predict PFS. Therefore the IISTH measures have potential clinical utility as they can be computed before treatment from non-invasive standard-of-care pre-operative CT images. Other known predictors of outcome, including surgical resection status [42, 43] and platinum resistance [44] can only be measured following treatment. The IISTH measures did not predict platinum resistance, but the MSKCC and TCIA datasets were dissimilar for known prognostic variables and platinum resistance information was missing for 37% of patients in the TCIA dataset. Average Haralick texture measures did not predict outcomes in either dataset.

We asked whether the cluDiss measure, which was an independent predictor of PFS was correlated with molecular features including enriched HALLMARK gene pathways and the tumour microenvironment. We found that the cluDiss measure showed a weak negative correlation to WNT/Beta-Catenin signaling enrichment and stromal cell types. Also, the enrichment of WNT/Beta-Catening signaling decreased with increasing cluster texture heterogeneity. These results, though preliminary due to dataset limitations, suggest that decreased textural heterogeneity between tumour sites may be associated with stronger WNT/Beta-Catenin signaling enrichment which in turn is associated with immune exclusion [41].

We performed exploratory analyses between IISTH measures and known putative prognostic genetic alterations. The ultra-high IISTH clusters from MSKCC had higher relative prevalence of patients with amplifications of *CCNE1* compared to low IISTH clusters, but did not reach statistical significance owing to the small sample size. Larger study populations are required to further explore radiogenomic hypotheses.

The IISTH measures were independent of scanner manufacturer. This is probably because IISTH measures variations in the texture dissimilarities between sub-regions rather than voxel-based texture values. They are therefore “meta” features that model the relative textural differences between the tumor sites and are robust to differences in texture values that may be different between scanner manufacturers. Reproducible estimates such as IISTH are essential for translating radiomics analysis for multi-centre trials.

Our study has several limitations. There were substantial differences between the MSKCC and TCIA datasets which prevent use of a training-validation set to establish clear cut points for clinical use. This underscores the difficulties of obtaining paired imaging and genomic data from multiple institutions to establish representative populations for study. Similar challenges for establishing cut points for predicting outcomes after checkpoint-blockage treatment have been observed where better outcomes are observed in individual datasets based on mutational load but no consistent cut-off has been identified [45–48].

We tried to overcome the limitation of heterogeneous datasets by employing unsupervised clustering (as opposed to supervised classification) to group the data and study association with outcomes. Specifically, we tested whether both datasets revealed the same trend in the association of the textural tumor heterogeneity with outcomes. The reported results are presented without adjusting for multiple comparisons as this is a feasibility study. Finally, examination of the correlations between IISTH measures and specific gene copy number alterations would ideally require a much larger cohort. However, our study represents the largest dataset for which both CT images and genomic information are publically available. Building larger data sets will provide a unique opportunity to probe correlations between radiomics and genomics and provide insights into the potential role of radiogenomics in predicting outcome in HGSOC.

## Conclusion

We developed quantitative, non-invasive texture-based measures to quantify intra- and inter-site tumor heterogeneity (IISTH) within patients with HGSOC and showed that these measures can be used to predict outcome across multiple institutions. Non-invasive imaging biomarkers such as IISTH can be computed from standard-of-care CT imaging requiring no additional tests for patients. These measures if validated on larger cohorts have the potential to identify high-risk patients early in their clinical course, thereby providing the opportunity to triage such patients to clinical trials that improve precision medicine for women with HGSOC.

## Supporting information

**S1 Methods** Dissimilarity matrix methods.

**S1 Fig.**
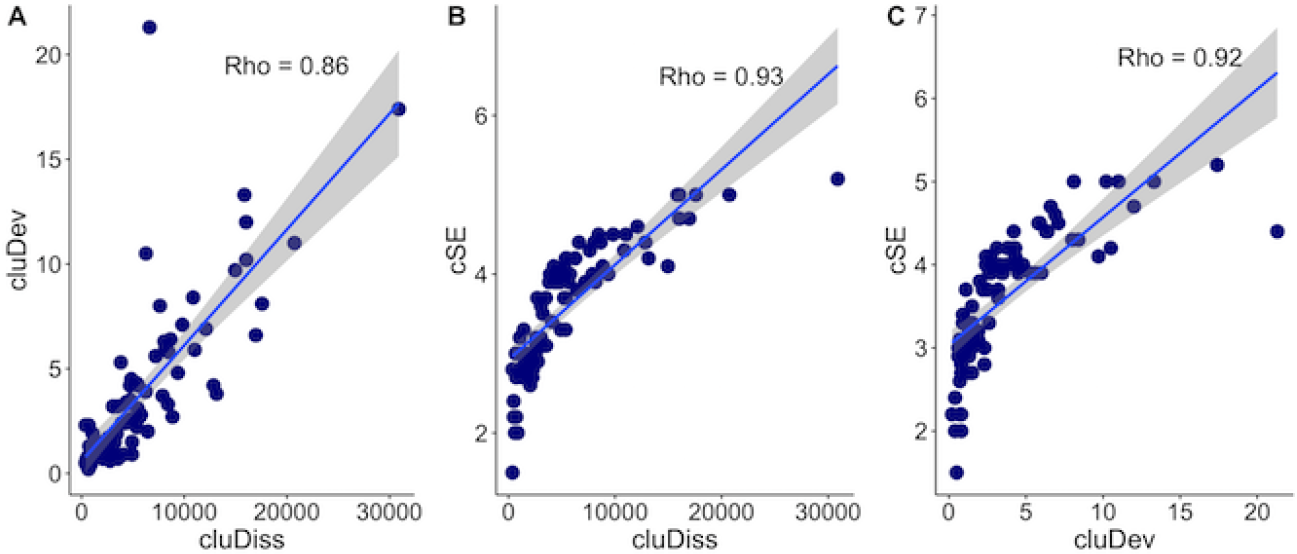
Spearman Rank correlation between the IISTH heterogeneity measures.

**S2 Fig.**
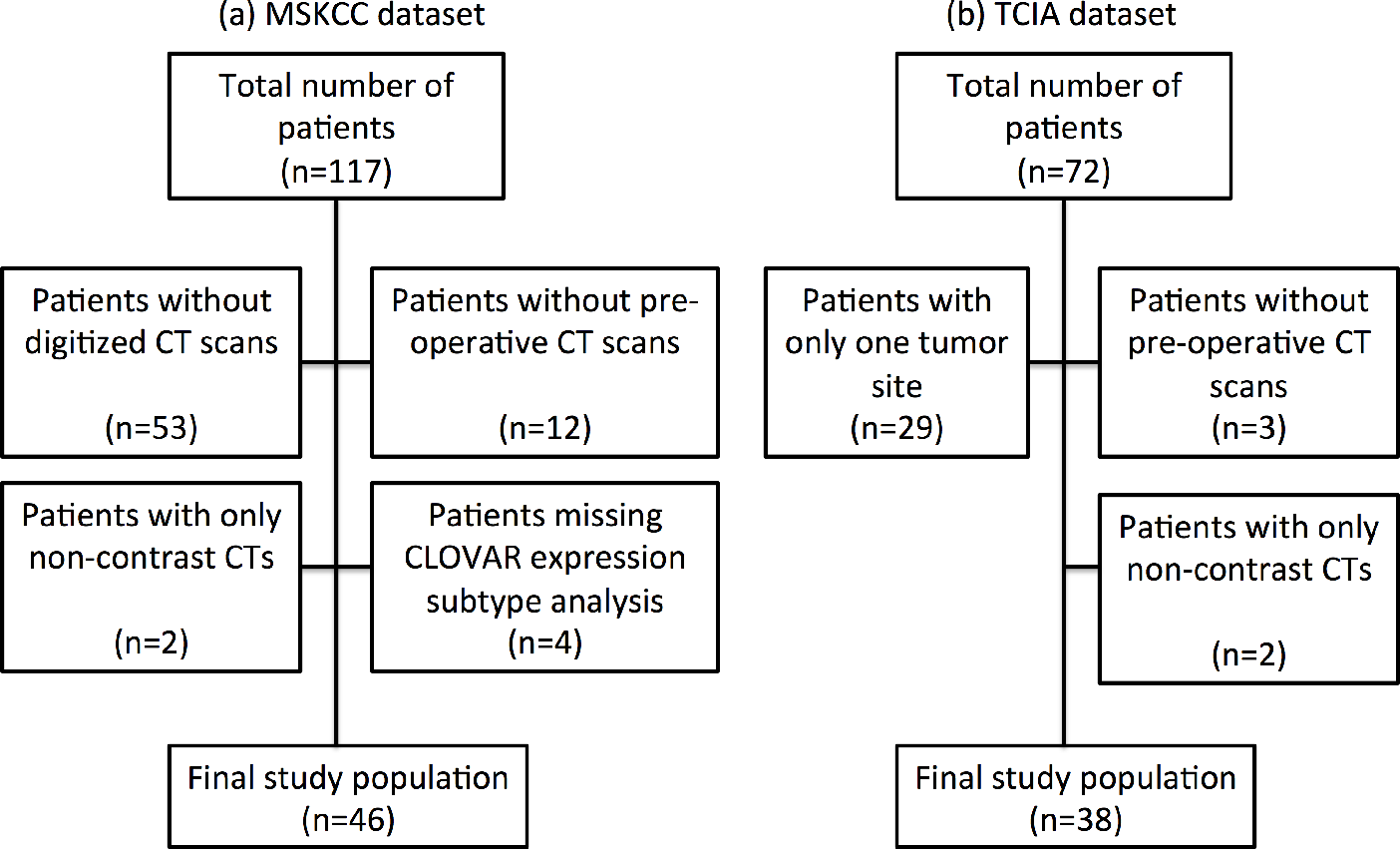
Flowchart of patient selection.

**S3 Fig.**
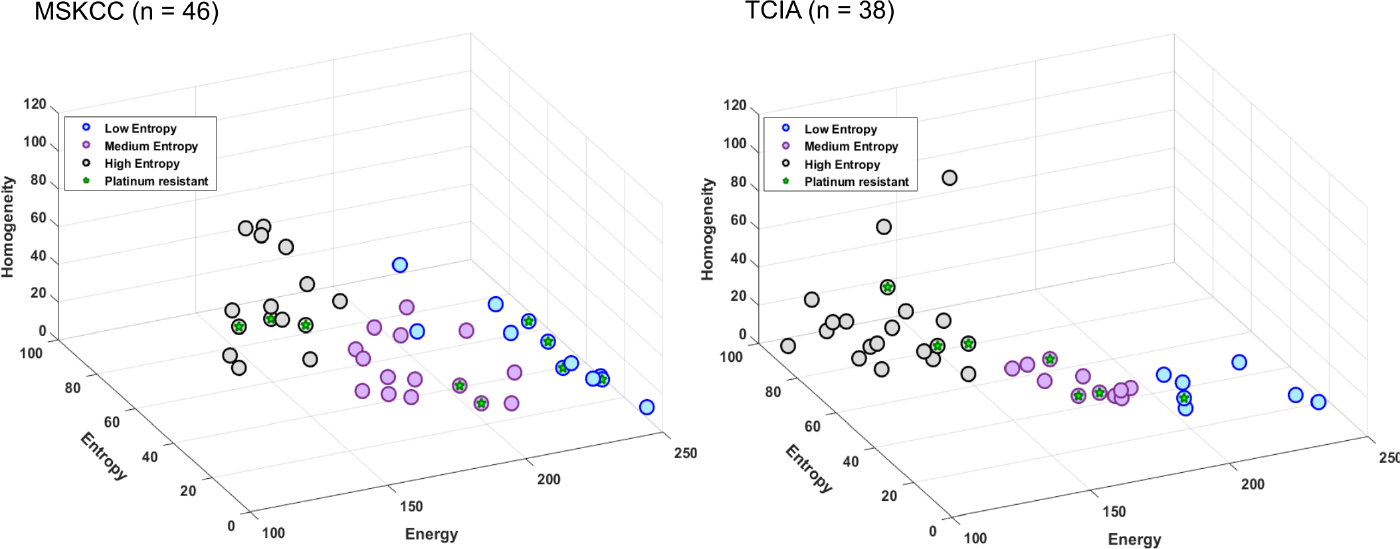
Results of unsupervised clustering using Haralick textures.

**S4 Fig.**
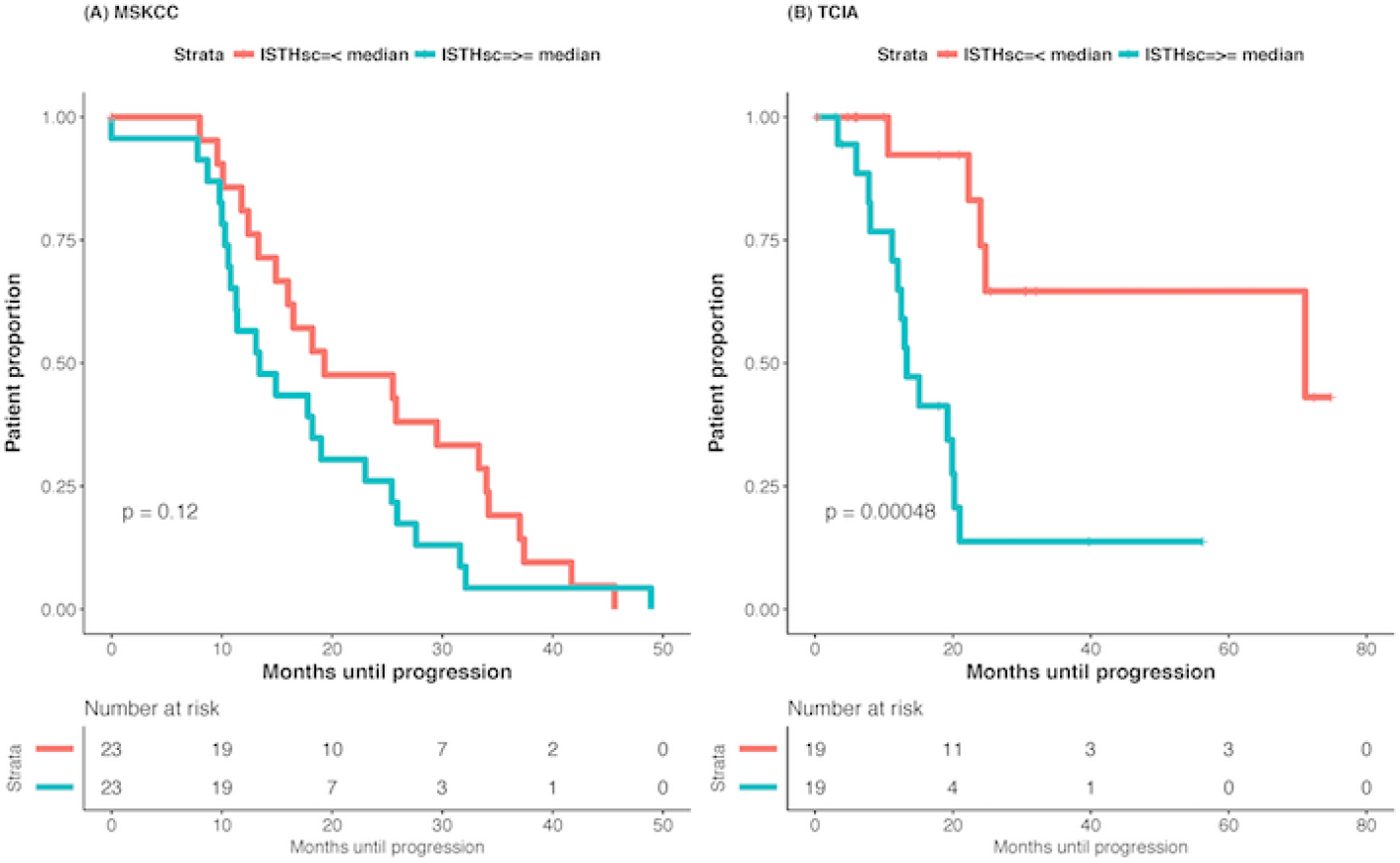
Kaplan-Meier curves computed using median values of the composite IISTH scores (MSKCC median = −0.022; TCIA median = −0.14). The IISTH scores were computed through factor analysis of the three IISTH measures consisting of cSE, cluDev, and cluDiss.

**S5 Fig.**
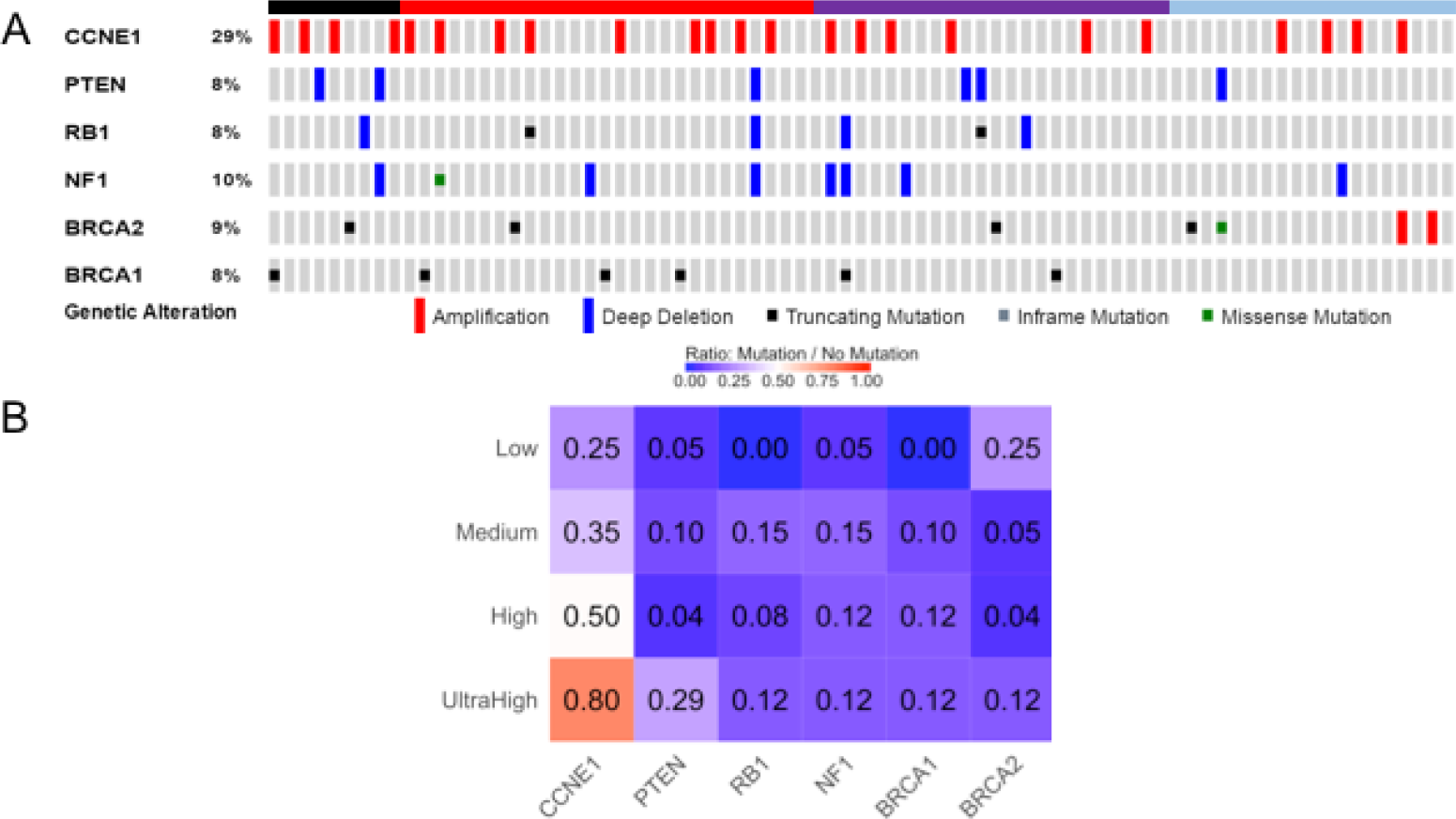
Prevalence of mutations in the IISTH clusters computed from TCIA and MSKCC datasets. A: shows the various gene mutations across all patients, organized by cluster heterogeneity. B: shows the ratio of mutation vs. absence of mutations within the clusters.

**S1 Table** Difference in Haralick texture and IISTH measures between MSKCC and TCIA dataset.

**S2 Table** Univariable (unadjusted) and multivariable (adjusted) analysis of pre-treatment values as predictors for overall survival (OS) for MSKCC and TCIA dataset. Multivariable analysis was adjusted for age, stage, and volume.

**S3 Table** Distribution of clinical characteristics in IISTH clustered groups for MSKCC and TCIA datasets.

**S4 Table** Distribution of clinical characteristics in Haralick textures-based clustered groups for MSKCC and TCIA datasets.

**S5 Table** Spearman rank correlation coefficient between CT texture-based measures and tumor burden.

## Acknowledgements

This research was partially supported by the MSK Cancer Center Support Grant/Core Grant (P30 CA008748) (but had no role in study design, data collection and analysis, decision to publish, or preparation of the manuscript) as well as the The Mark Foundation for Cancer Research and Cancer Research UK Cambridge Centre [C9685/A25177].

## Material for Supporting information

## S1_Methods

### Computing inter-site tumor heterogeneity (IISTH) measures

The steps in computing the IISTH measures are summarized in Fig 1 together with the method used in each step. Details of the individual steps are discussed in the following subsections.

**Fig 1.**
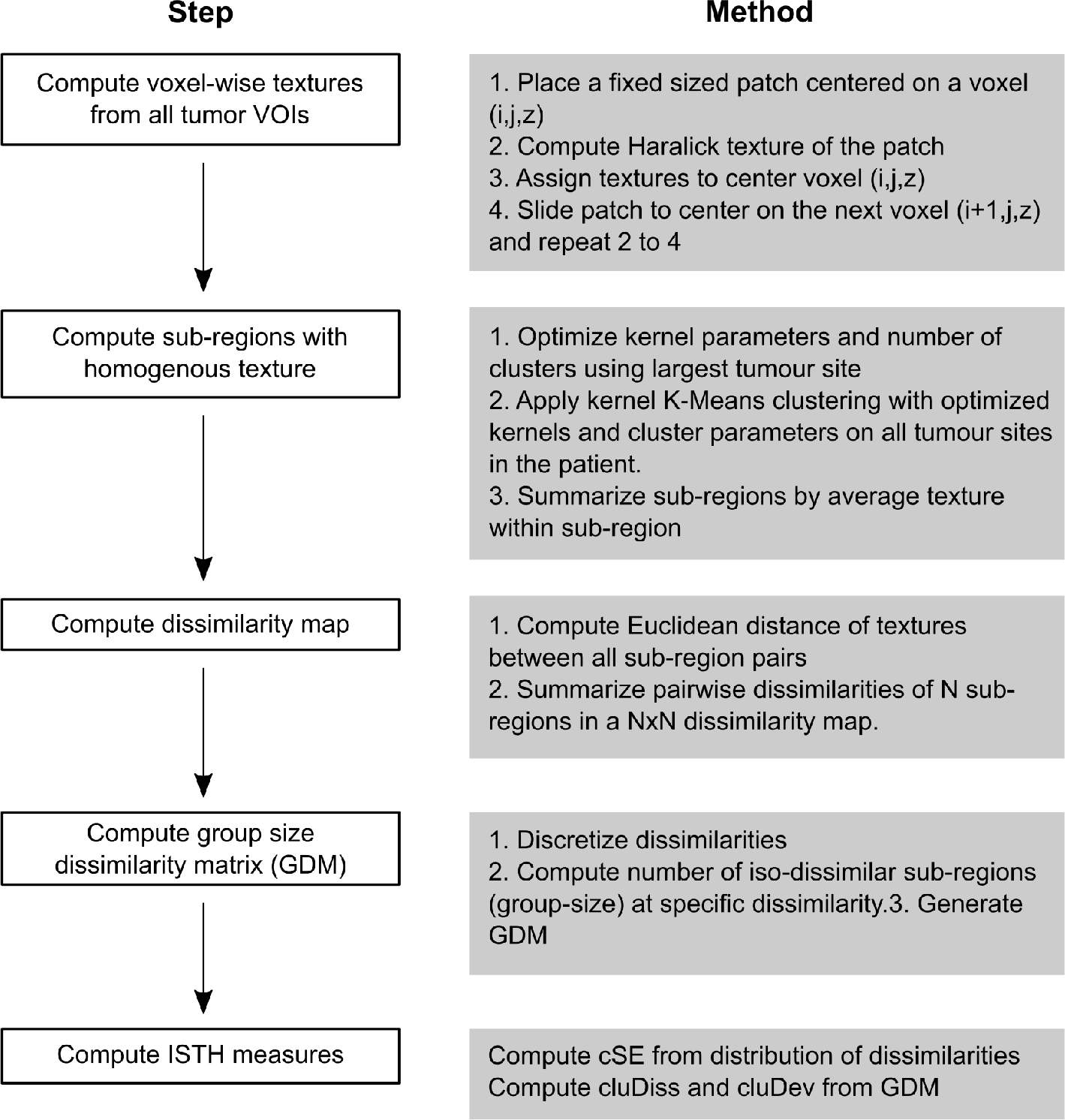
Steps in generating IISTH measures.

### Extracting sub-regions of homogeneous texture

The first step in the IISTH measure computation is the extraction of voxel-wise textures by sliding fixed sized patches across the whole image. The texture measures computed using Haralick textures is assigned to the voxel in the centre of that patch. Next, sub-regions of homogeneous texture were computed by clustering of the voxel-wise textures using kernel K-means method. The appropriate number of clusters and the kernel parameters (mean *μ*, standard deviation *Σ*) were computed from the largest tumor site using expectation maximization (EM) algorithm. Akaike information criterion (AIC) was used to select the best number of clusters. The number of analysed clusters ranged from two to five. The learned kernel parameters were then used to produce sub-regions in the remaining tumor sites. As a result of this step, all the tumor sites were divided into distinct sub-regions. The subregions were described using the mean of the individual texture measures.

### Dissimilarity matrix methods

The dissimilarity between two sub-regions is computed using the Euclidean distance of the sub-region textures. Pair-wise sub-region dissimilarities are represented using a dissimilarity matrix *D* where the cells *D*_*i*, *j*_ correspond to dissimilarity between sub-regions *i* and *j*. The elements of the dissimilarity matrix are organized by numerical labels of the sub-regions extracted within each anatomical structure such that the total number of elements within the dissimilarity matrix equals the total number of all sub-regions. Heterogeneous dissimilarity will result when sub-regions have highly distinct textures. Conversely, homogeneous textural differences between sub-regions will give rise to a homogeneously appearing dissimilarity map.

Pairwise sub-region dissimilarities at the patient level are discretized into ten bins of increasing normalized dissimilarity (ranging from 0 to 1). The IISTH metric cSE is computed from the frequencies of pairwise sub-region dissimilarities as:

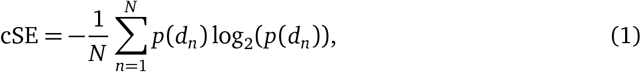

where *N* are the number of bins, *p*(*d*) corresponds to the normalized dissimilarity *d*. The dissimilarities are normalized using the z-score prior to computation of cSE.

### Group Dissimilarity Matrix

The ISTH measures are computed from the dissimilarity map by converting it into a group dissimilarity matrix (GDM). GDM is a two dimensional histogram computed from the dissimilarity matrix *D* that captures the prevalence of dissimilarities shared by different number of sub-regions. The elements of the GDM are the number of discrete dissimilarities and the number of sub-region pairs sharing those dissimilarities. GDM is constructed similar to zone-level size matrices [49] and captures the spread of dissimilarities. Texturally identical sub-regions will give rise to a single peak at the largest group size and lowest dissimilarity value of the GDM. On the other hand, presence of highly distinct textures in the sub-regions of various anatomic structures will give rise to uniform distribution of the dissimilarities at lower group sizes. The GDM matrix was always ordered in the same way by the individual metastatic sites starting from the primary site with the subsequent sites included in anti-clockwise order.

The measures cluDev and cluDiss are computed from the GDM. Cluster standard deviation (cluDev) is computed as the standard deviation of the GDM. Cluster dissimilarity (cluDiss) is computed as:

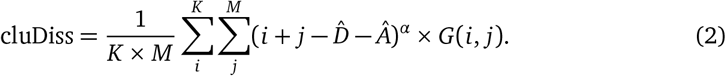

where *K*, *M* are the number of discrete dissimilarity and group size levels, 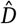 is the mean dissimilarity, 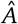 is the mean group size, and *G* is the GDM, with *α* = 4. Varying the value of *α* alters the emphasis on the mean dissimilarities. The indices *i* and *j* emphasize larger dissimilarities and larger group sizes thereby, resulting in large values of cluDiss in the presence of many texturally distinct sub-regions from all other sub-regions as shown in Fig 2B. On the other hand, presence of large groups with distinct dissimilarity will result in small cluDiss values as shown in Fig 2A.

**Fig 2.**
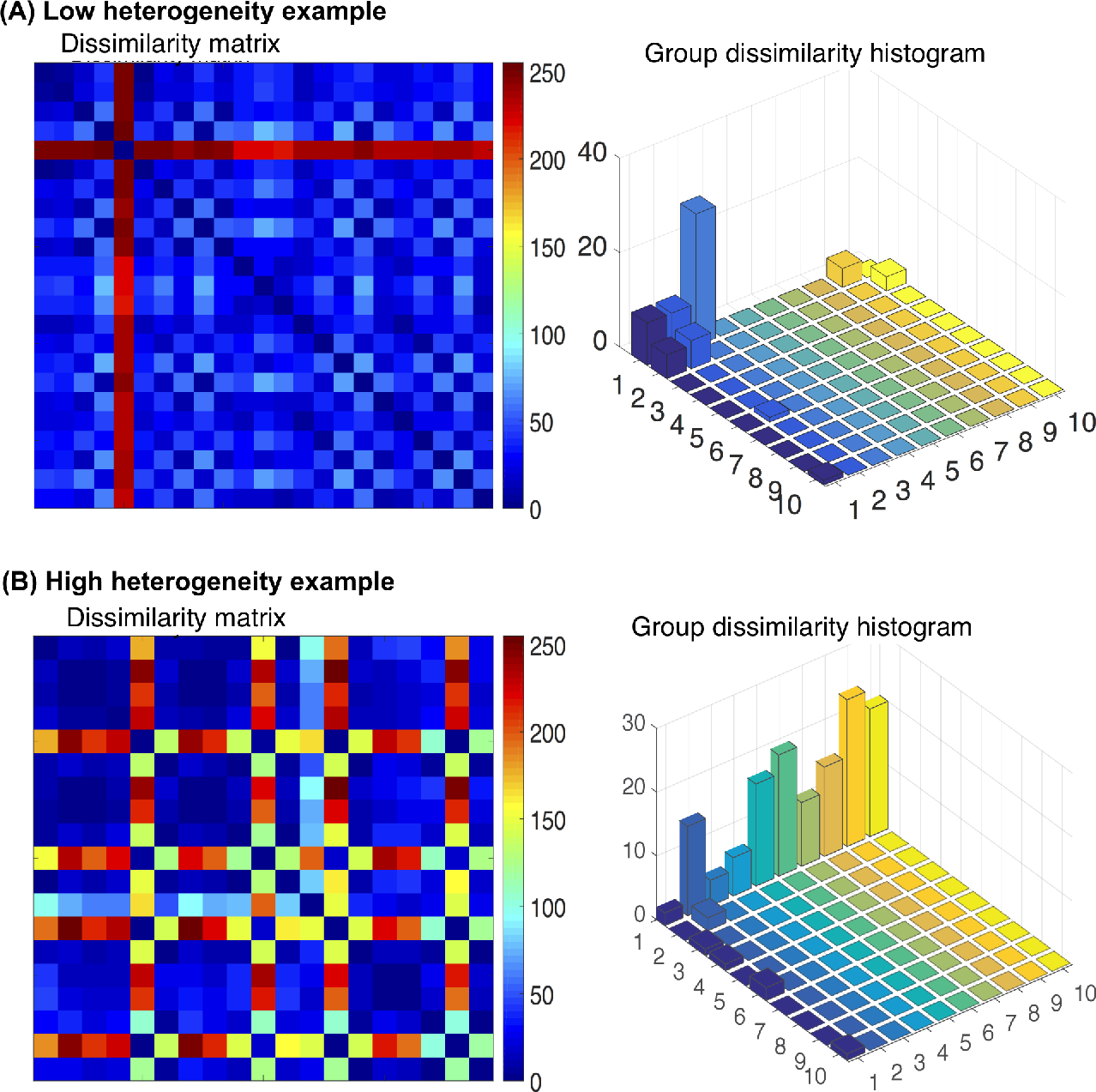
Examples of low and high inter-site texture heterogeneity. Presence of homogeneous distribution of dissimilarities across sub-regions results in few distinct peaks in the corresponding GDM matrix (A). Presence of many sub-regions with distinct texture results in uniformly distributed dissimilarities in the GDM and high heterogeneity (B).

**S1_Table.**
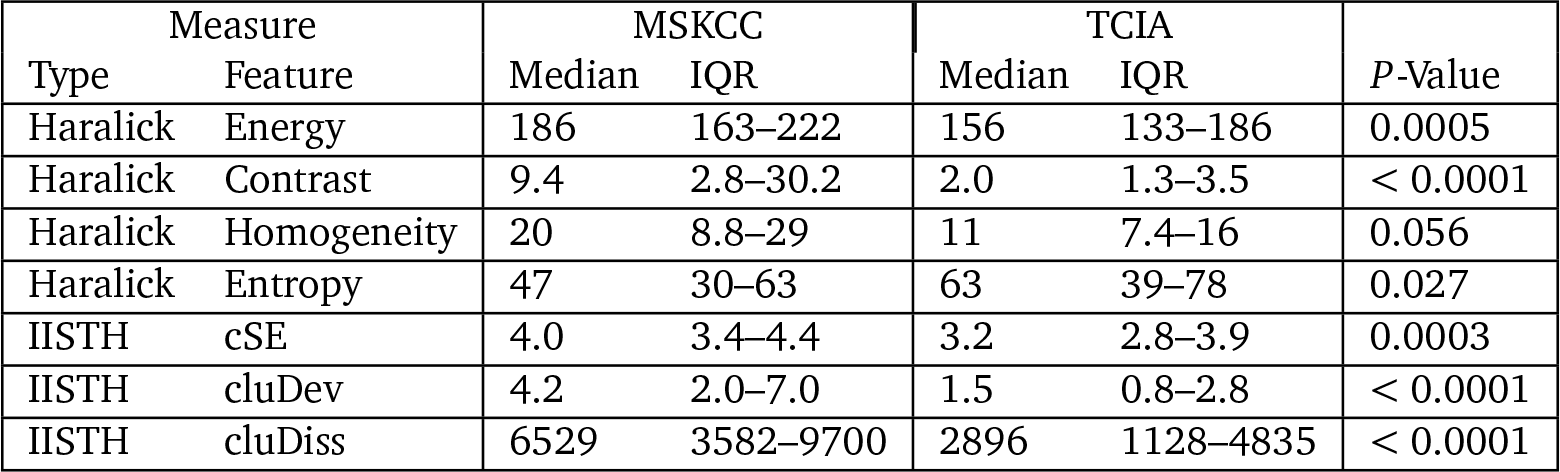
Differences in average texture (Haralick) and IISTH measures between MSKCC and TCIA datasets.

**S2_Table.** Univariable (unadjusted) and multivariable (adjusted) analysis of pre-treatment values as predictors for overall survival (OS) for TCIA dataset. Multivariable Cox hazard regression was adjusted for age, stage, resection status, and volume.

| Variable | MSKCC |  |  |  |  |  | TCIA |  |  |  |  |  |
| --- | --- | --- | --- | --- | --- | --- | --- | --- | --- | --- | --- | --- |
|  | Univariable |  |  | Multivariable |  |  | Univariable |  |  | Multivariable |  |  |
|  | HR | CI | P-Value | HR | CI | P-Value | HR | CI | P-Value | HR | CI | P-Value |
| cSE | 1.50 | 0.89–2.50 | 0.130 | 1.10 | 0.46–2.70 | 0.81 | 1.10 | 0.60–1.90 | 0.810 | 2.80 | 0.66–12.3 | 0.161 |
| cluDiss | 1.07 | 1.02–1.10 | 0.013 | 1.00 | 1.00–1.00 | 0.117 | 0.93 | 0.81–1.10 | 0.277 | 1.00 | 1.00–1.00 | 0.474 |
| cluDev | 1.03 | 0.96–1.10 | 0.40 | 0.97 | 0.86–1.10 | 0.691 | 0.87 | 0.62–1.20 | 0.41 | 0.91 | 0.45–1.90 | 0.806 |
| IISTH score | 1.50 | 0.63–3.50 | 0.365 | 1.50 | 0.63–3.50 | 0.365 | 0.99 | 0.67–1.48 | 0.98 | 2.00 | 0.65–5.80 | 0.23 |
| IISTH cluster (3,4/1,2) | 3.47 | 1.55–7.80 | 0.002 | 4.00 | 1.32–12.4 | 0.014 | 0.84 | 0.36–2.00 | 0.693 | 2.10 | 0.52–8.60 | 0.298 |
| Energy | 1.00 | 0.99–1.00 | 0.490 | 1.00 | 0.99–1.00 | 0.669 | 1.00 | 0.99–1.00 | 0.780 | 0.99 | 0.98–1.0 | 0.121 |
| Entropy | 0.99 | 0.98–1.00 | 0.210 | 1.0 | 0.99–1.00 | 0.738 | 1.00 | 0.98–1.00 | 0.910 | 1.00 | 0.99–1.00 | 0.155 |
| Contrast | 0.98 | 0.96–1.00 | 0.090 | 0.99 | 0.96–1.00 | 0.16 | 1.03 | 1.00–1.10 | 0.024 | 1.00 | 0.99–1.10 | 0.11 |
| Homogeneity | 0.97 | 0.95–1.00 | 0.019 | 0.98 | 0.96–1.00 | 0.235 | 1.00 | 0.97–1.00 | 0.878 | 1.00 | 0.95–1.00 | 0.855 |
| Haralick cluster (3/4) | 0.78 | 0.30–2.00 | 0.610 | 1.20 | 0.41–3.60 | 0.726 | 0.95 | 0.34–2.60 | 0.920 | 3.00 | 0.77–11.60 | 0.112 |
| Haralick cluster (2/1) | 0.79 | 0.31–2.00 | 0.62 | 0.85 | 0.30–2.40 | 0.76 | 2.20 | 0.60–7.70 | 0.237 | 2.40 | 0.63–9.30 | 0.20 |
| Resection (sub-optimal) | 6.00 | 1.99–18.0 | 0.001 | 3.90 | 1.15–13.3 | 0.029 | 7.2 | 1.40–37.0 | 0.018 | 5.70 | 1.09–29.6 | 0.039 |
| Resection (optimal) | 1.80 | 0.66–4.70 | 0.261 | 1.40 | 0.52–3.80 | 0.502 | 4.60 | 1.03–20.8 | 0.046 | 5.10 | 1.10–23.8 | 0.038 |
| Stage (IV/III,II) | 1.20 | 0.52–2.60 | 0.731 | 1.41 | 0.53–3.80 | 0.488 | 3.0 | 0.93–9.6 | 0.066 | 2.33 | 0.78–6.9 | 0.128 |
| Age (years) | 1.10 | 1.01–1.10 | 0.021 | 1.00 | 1.00–1.10 | 0.055 | 1.00 | 0.99–1.10 | 0.156 | 1.00 | 0.98–1.10 | 0.265 |
| Volume (cm<sup>3</sup>) | 1.00 | 1.00–1.00 | 0.04 | 1.00 | 0.99–1.00 | 0.195 | 0.297 | 0.99–1.00 | 0.250 | 1.00 | 1.00–1.00 | 0.290 |
| Sites | 1.18 | 1.03–1.3 | 0.017 | 1.10 | 0.90–1.30 | 0.421 | 0.92 | 0.69–1.20 | 0.607 | 0.92 | 0.61–1.40 | 0.706 |

Haralick clusters are coded 1–4 for low, medium, high and ultra-high heterogeneity.
For continuous measures, HR > 1, an increase in the value is associated with higher risk or number of events and decreased OS. For binary variables, the HR listed is the first option and HR=1 is the second option. HR < 1 indicates lowering of risk with the same option.
CI - confidence interval HR - hazard ratio.
Cox analysis for cluDiss as a continuous measure was computed after computing square root of cluDiss to achieve reasonable data scaling.

**S3_Table.** Distribution of clinical characteristics in IISTH clustered groups for MSKCC and TCIA datasets.

| Patient<br>Characteristic | MSKCC |  |  |  | TCIA |  |  |  |
| --- | --- | --- | --- | --- | --- | --- | --- | --- |
|  | Low | Medium | High | Ultra-high | Low | Medium | High | Ultra-high |
| Number of cases, *n* (%) | 9 (20) | 15 (33) | 14 (30) | 8 (17) | 13 (34) | 10 (26) | 13 (34) | 2 (5) |
| PFS median mos (IQR) | 33 (10–37) | 19 (13–28) | 13 (11–21) | 14 (9–20) | 22 (6–32) | 15 (7–29) | 13 (12–20) | 11 (7–15) |
| OS median mos (IQR) | 76 (59–109) | 69 (54–76) | 47 (33–63) | 50 (25–77) | 32 (6–60) | 18 (7–29) | 36 (27–40) | 80 (60–100) |
| Platinum resistant, *n* (%) | 1 (11) | 1 (7) | 5 (36) | 2 (25) | 1 (8) | 2 (20) | 3 (23) | 1 (50) |
| Platinum sensitive, *n* (%) | 8 (23) | 13 (37) | 9 (26) | 5 (14) | 4 (24) | 3 (18) | 9 (53) | 1 (6) |
| Complete resection, *n* (%) | 7 (78) | 5 (33.3) | 2 (14.3) | 1 (12.5) | 2 (15) | 2 (20) | 3 (23) | 1 (50) |
| Optimal resection, *n* (%) | 1 (11) | 8 (53.3) | 9 (64.3) | 5 (62.5) | 7 (54) | 2 (20) | 7 (54) | 0 (0) |
| Suboptimal resection, *n* (%) | 1 (11) | 2 (13.3) | 3 (21.4) | 2 (25) | 3 (23) | 5 (50) | 3 (23) | 1 (50) |

Survival and follow-up times are reported in months.

**S4_Table.** Distribution of clinical characteristics in Haralick textures-based clustered groups for MSKCC and TCIA datasets.

| Patient Characteristic | MSKCC |  |  | TCIA |  |  |
| --- | --- | --- | --- | --- | --- | --- |
|  | Low | Medium | High | Low | Medium | High |
| Number of cases, *n* (%) | 15 (33) | 15 (33) | 16 (35) | 7 (18) | 11 (29) | 20 (53) |
| PFS median mos (IQR) | 16 (12–26) | 19 (12–29) | 13 (10–31) | 22 (17–52) | 10 (5–19) | 16 (10–24) |
| OS median mos (IQR) | 57 (42–68) | 69 (43–83) | 56 (47–85) | 40 (35–63) | 18 (5–28) | 32 (17–45) |
| Platinum resistant *n* (%) | 4 (27) | 2 (13) | 3 (19) | 1 (14) | 3 (27) | 3 (15) |
| Longer PFS *n* (%) | 0 (0) | 1 (7) | 1 (6) | 4 (57) | 8 (73) | 7 (35) |
| Complete resection *n* (%) | 2 (13) | 6 (40) | 7 (44) | 0 (0) | 1 (9) | 7 (35) |
| Optimal resection *n* (%) | 9 (60) | 6 (40) | 8 (50) | 5 (71) | 5 (45.5) | 6 (30) |
| Suboptimal resection *n* (%) | 4 (27) | 3 (20) | 1 (6) | 2 (29) | 5 (45.5) | 5 (25) |

**S5_Table.** Spearman rank correlation coefficient between CT texture-based measures and tumor burden.

| Measure |  | Tumor volume (cm<sup>3</sup>) |  | Sites (N) |  |
| --- | --- | --- | --- | --- | --- |
| Type | Feature | $\rho$ | *P*-value | $\rho$ | *P*-value |
| IISTH | cSE | 0.016 | 0.883 | 0.846 | < 0.001 |
| IISTH | cluDiss | 0.037 | 0.735 | 0.866 | < 0.001 |
| IISTH | cluDev | 0.012 | 0.914 | 0.889 | < 0.001 |
| Haralick | Energy | −0.039 | 0.727 | 0.432 | < 0.001 |
| Haralick | Entropy | −0.157 | 0.153 | −0.436 | < 0.001 |
| Haralick | Contrast | −0.417 | < 0.001 | 0.194 | 0.077 |
| Haralick | Homogeneity | −0.412 | < 0.001 | −0.114 | 0.304 |

